# Chemosensory proteins in the CSP4 clade evolved as plant immunity suppressors before two suborders of plant-feeding hemipteran insects diverged

**DOI:** 10.1101/173278

**Authors:** Claire Drurey, Thomas C. Mathers, David C. Prince, Christine Wilson, Carlos Caceres-Moreno, Sam T. Mugford, Saskia A. Hogenhout

## Abstract

Chemosensory proteins (CSPs) are small globular proteins with hydrophobic binding pockets that have a role in detection of chemicals, regulation of development and growth and host seeking behaviour and feeding of arthropods. Here, we show that a CSP has evolved to modulate plant immune responses. Firstly, we found that the green peach aphid *Myzus persicae* CSP Mp10, which is delivered into the cytoplasm of plant cells, suppresses the reactive oxygen species (ROS) bursts to both aphid and bacterial elicitors in *Arabidopsis thaliana* and *Nicotiana benthamiana*. Aphid RNA interference studies demonstrated that Mp10 modulates the first layer of the plant defence response, specifically the BAK1 pathway. We identified Mp10 homologs in diverse plant-sucking insect species, including aphids, whiteflies, psyllids and leafhoppers, but not in other insect species, including blood-feeding hemipteran insects. We found that Mp10 homologs from other splant-sucking insect species are also capable of suppressing plant ROS. Together, these data and phylogenetic analyses provides evidence that an ancestral Mp10-like sequence acquired plant ROS suppression activity before the divergence of plant-sucking insect species over 250 million years ago.

**Significance:** Aphids, whiteflies, psyllids, leafhoppers and planthoppers are plant-sucking insects of the order Hemiptera that cause dramatic crop losses via direct feeding damage and vectoring of plant pathogens. Chemosensory proteins (CSPs) regulate behavioural and developmental processes in arthropods. Here we show that the CSP Mp10 of the green peach aphid *Myzus persicae* is an effector that suppresses plant reactive oxygen species (ROS) bursts and the first layer of plant defence responses. Surprisingly, Mp10 homologs are present in diverse plant-feeding hemipteran species, but not blood-feeding ones. An ancestral Mp10-like sequence most likely acquired ROS suppression activity before the divergence of plant-sucking insect species 250 million years ago.

## Introduction

Chemosensory proteins (CSPs) are soluble and stable proteins consisting of 6 alpha-helices stabilized by two disulphide bonds and a central channel with the capacity to bind small hydrophobic molecules, such as plant volatiles and insect pheromones (1, 2). These proteins are often highly expressed in the olfactory and gustatory organs of insects in which they play a role in the sensing of the external environment by carrying volatiles and pheromones to neurons of chemosensilla, leading to downstream behavioural and developmental processes (3, 4). For example, CSPs expressed in antenna regulate the transition from solitary to migratory phases of migratory locusts (5), female host-seeking behaviour of tsetse flies (6) and nest mate recognition of ants (2).

However, CSPs are also found in tissues that do not have chemosensory functions. For example, CSPs regulate embryo development in honey bees (7), limb regeneration in cockroaches (8) and immune protection against insecticides (9). Three CSPs are expressed in the midgut of the lepidopteran insect *Spodoptera litura*, and are differentially expressed in response to diets and bind non-volatile chemicals typically found in plants (10). A CSP highly abundant in the lumen of mouthparts that is evenly distributed along the length of the proboscis of *Helicoverpa* species does not seem involved in chemodetection at all, but acts as a lubricant to facilitate acquisition of sugar solution from plants via sucking (11), though orthologous proteins in the proboscis of four other lepidopterans bind the plant compound ß-carotene (12). Interestingly, the same CSPs are also highly expressed in the eyes of the lepidopterans and bind visual pigments abundant in eyes that are related to ß-carotene (12). CSPs may therefore be expressed in multiple tissues where they regulate a variety of processes in insects often, but not always, upon binding small molecules.

Aphids have about 10 CSP genes, and similar numbers were identified in related plant-sucking insects of the order Hemiptera, such as the whitefly *Bemisia tabaci* and psyllid *Diaphorina citri* (13–16). Aphid CSPs were previously known as OS-D-like proteins (17). OS-D1 and OS-D2 transcripts and proteins were detected in antennae, legs and heads suggesting a chemosensory role (17). However, OS-D2 did not bind any of 28 compounds, including (E)-ß-farnesene and related repellents and several other volatile plant compounds (17). Unexpectedly, a screen developed for identification of virulence proteins (effectors) in the saliva of the green peach aphid *Myzus persicae* identified OS-D2, named Mp10 in the screen, as a suppressor of the plant reactive oxygen species (ROS) burst, which is part of the plant defence response (18). Mp10/OS-D2 (henceforth referred to as Mp10) also modulates other plant defence responses (19). Moreover, the protein was detected in the cytoplasm of plant mesophyll cells adjacent to sucking-sucking mouthparts (stylets) of aphids, indicating that the aphid stylets deposit Mp10 into these cells during navigation to the plant vascular tissue for long-term feeding (20). Taken together, these data suggest a role of Mp10 in plants. However, so far, there is no evidence that CSPs have functions beyond arthropods. Hence, we investigated the role of Mp10 further.

Here we show that the *M. persicae* CSP Mp10 modulates the first layer of the plant defence response that is induced to aphid attack. This ROS burst suppression activity is shared among orthologous proteins of Mp10 in diverse plant-feeding hemipteran insects, including aphids and whiteflies (Sternorrhyncha) and leafhoppers (Auchenorrhyncha) (21). It is likely that an ancestral Mp10-like sequence acquired activity to suppress plant immunity before the divergence of plant-feeding hemipterans.

## Results

### Mp10 blocks plant ROS bursts

Plant defence responses are induced upon detection of pathogen or pest elicitors, such as pathogen/microbe-associated molecular patterns (PAMPs/MAMPs), by cell-surface receptors (22, 23). For example, the 22-amino acid sequence of the conserved N-terminal part of bacterial flagellin (flg22) is a well-characterized PAMP that binds the plant cell-surface receptor flagellin-sensitive 2 (FLS2) leading to interaction with the co-receptor BRASSINOSTEROID INSENSTIVE 1-associated receptor kinase 1 (BAK1) and initiation of plant defence responses, including a ROS burst, in *Arabidopsis thaliana* and *Nicotiana benthamiana* (24). Elicitors identified in whole extracts of aphids also induce defence responses, including ROS bursts, in a BAK1-dependent manner (25, 26). Hence, we first investigated if Mp10 suppresses ROS burst induced to flg22 and aphid elicitors in *A. thaliana* and *N. benthamiana*, which are readily colonized by *M. persicae* clone O (27). Efforts to stably express Mp10 in *A. thaliana* were unsuccessful, perhaps because high concentrations of Mp10 are toxic to plants, in agreement with this protein inducing severe chlorosis and plant defence responses (18, 19). However, we found that GFP and GFP-Mp10 can transiently be produced in *N. benthamiana* leaves via *Agrobacterium*-mediated infiltration (18). Here we show that this method allows the transient production of these three proteins in *A. thaliana* leaves (Supplementary file, Fig. 1). GFP-Mp10 suppressed the ROS burst to both aphid elicitors and flg22 in *A. thaliana* and *N. benthamiana*, whereas GFP alone did not (Fig. 1; Supplementary file, Fig. 2). Therefore, Mp10 supresses ROS bursts to diverse elicitors in at least two distantly related plant species.

**Figure 1:**
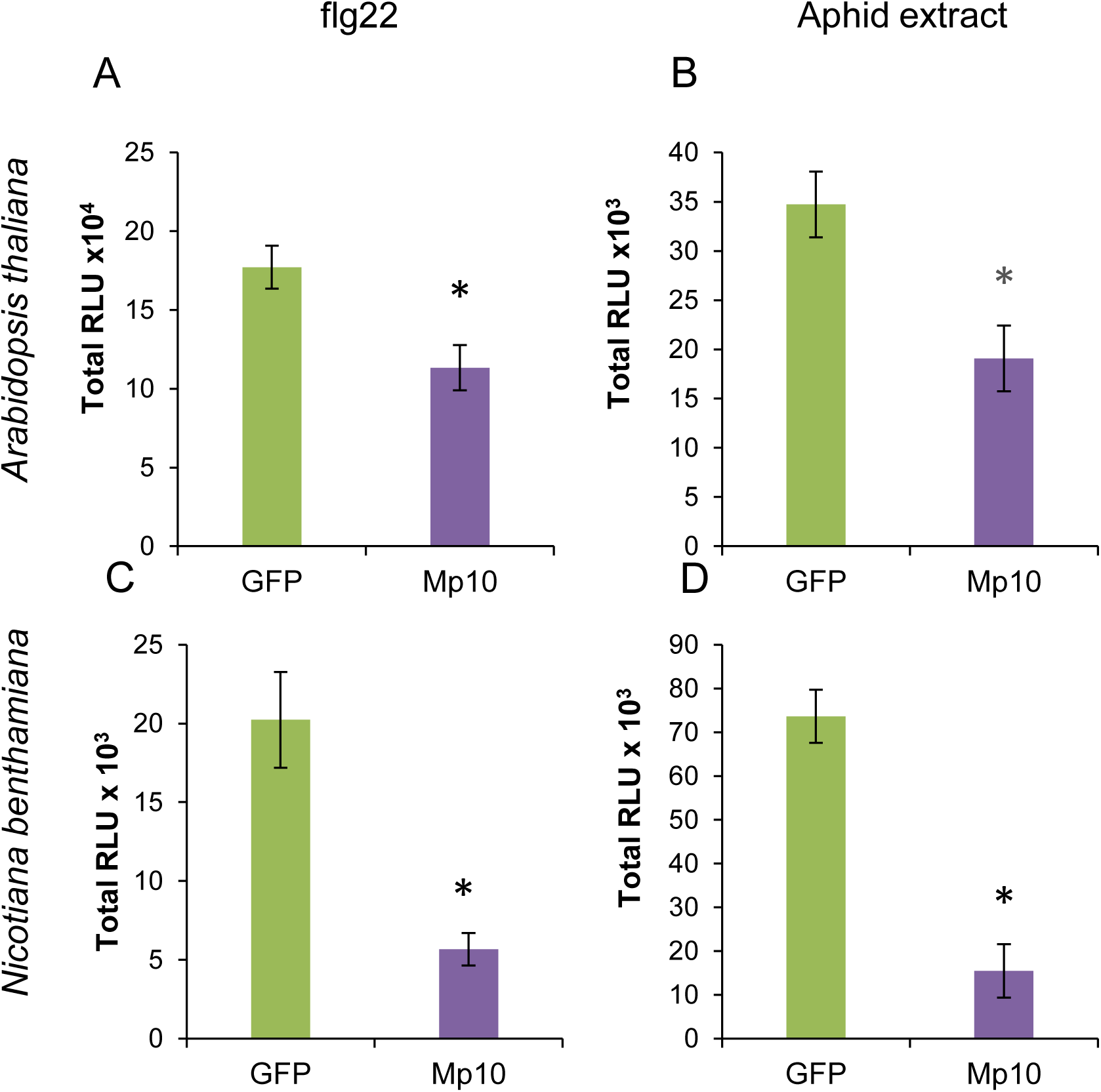
Mp10 suppresses elicitor-induced ROS bursts in *A. thaliana* and *N. benthamiana.* Total ROS bursts were measured as relative light units (RLU) in luminol-based assays of *A. thaliana* (A, B) and *N. benthamiana* (C, D) leaves upon elicitation with flg22 (A, C) or aphid extract (B, D). The leaves transiently produced GFP-tagged Mp10 (GFP-Mp10) alongside a GFP control (Figure S1). Bars show mean ± SE of total RLUs measured over periods of 0-60 minutes (flg22), 0-320 minutes (*N. benthamiana*, aphid extract) or 0-600 minutes (*A*. *thaliana*, aphid extract) of three (A, B, C) or two (D) independent experiments (n=8 per experiment). Asterisks indicate significant differences to the GFP treatment (Student’s t-probability calculated within GLM at P <0.05).

### Mp10 is required for *M. persicae* colonisation of Arabidopsis in a BAK1-dependent manner

Given that BAK1 is required for the flg22-mediated ROS burst (24, 29) and is involved in plant defence to aphids (25, 26), we determined if Mp10 acts in the BAK1 pathway during aphid feeding. Transgenic plants producing double-stranded (ds)RNA corresponding to *M. persicae Mp10* (*dsMp10*) were generated for knock down of *Mp10* expression in *M. persicae* by plant-mediated RNA interference (RNAi) (30, 31) in both *A. thaliana* Col-0 wild type (WT) and *bak1* mutant backgrounds, alongside *dsGFP* and *dsRack1* transgenic plants as controls (30, 31). Rack1 is an essential regulator of many cellular functions (32). The expression levels of *Mp10* and *Rack1* were similarly reduced by about 40% in the RNAi aphids on the dsRNA WT and *bak1* plants compared to the *dsGFP*-exposed aphids (Fig. 2 B, C). On the dsRNA WT plants, *Rack1*-RNAi and *Mp10*-RNAi aphids produced about 20% less progeny compared to ds*GFP*-exposed aphids (Fig. 2A). On the dsRNA *bak1* plants however, the *Mp10*-RNAi aphids produced similar levels of progeny compared to the *dsGFP*-exposed aphids, and only the *Rack1*-RNAi aphids had a reduced fecundity of about 20%, similar to that of *Rack1*-RNAi aphids on WT Col-0 plants (Fig. 3A). Aphid survival rates were not different among the treatments on the dsRNA WT and *bak1* plants (Supplementary file, Fig. 3). The fecundity of *Mp10*-RNAi aphids is therefore reduced on *dsMp10* WT but not on *dsMp10 bak1* plants, suggesting that Mp10 acts in the BAK1 signalling pathway. In contrast, *Rack1*-RNAi aphids had reduced fecundity on both WT and *bak1* plants, in agreement with Rack1 being involved in the regulation of cell proliferation, growth and movement in animals and having no known roles in plant-insect interactions (32).

**Figure 2:**
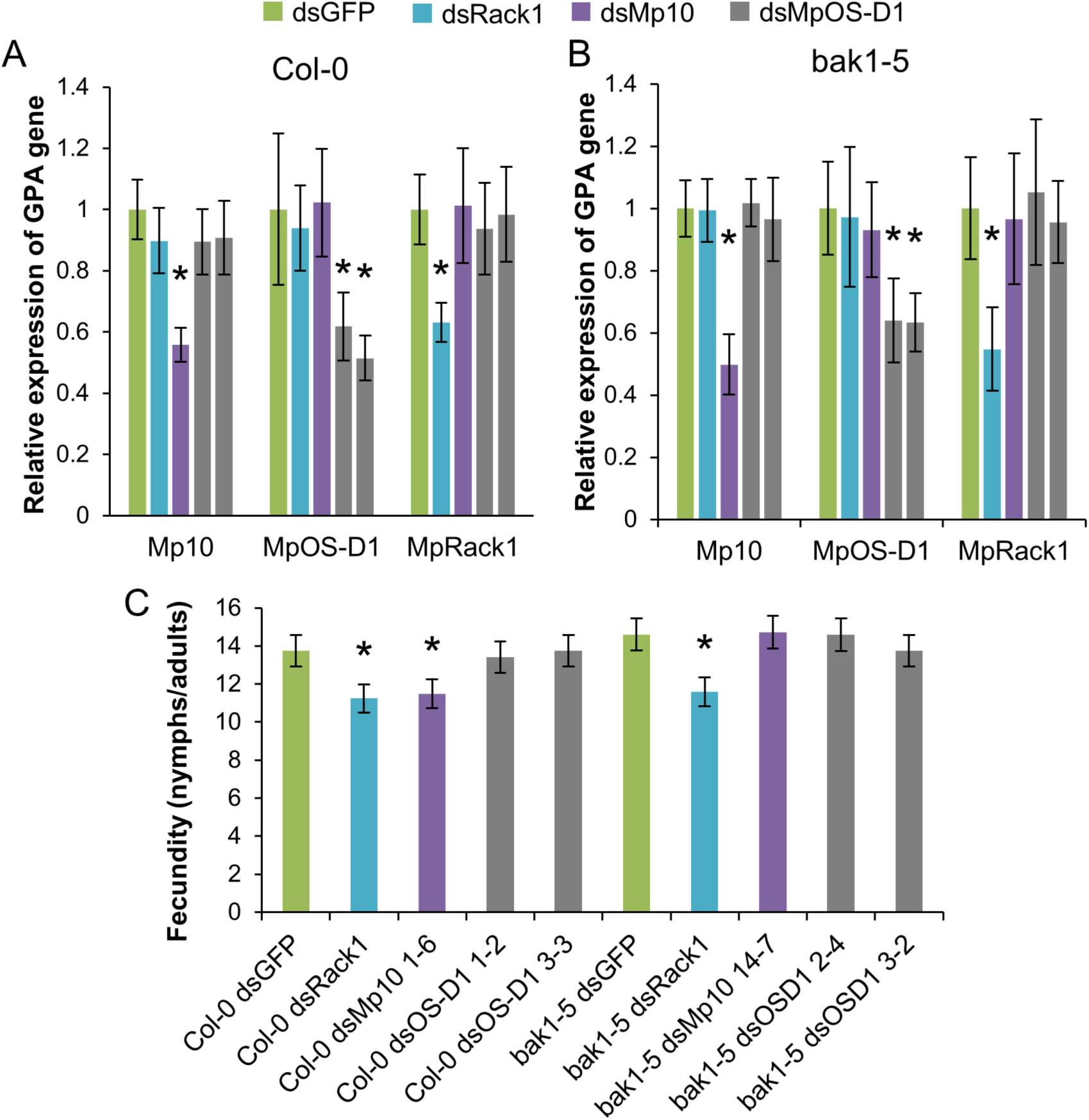
Knock down of *Mp10* expression reduces aphid reproduction on *A. thaliana* wild type, but not on *bak1-5* mutants. (A, B) Relative expression levels of Mp10, MpOS-D1 and MpRack1 genes in M. persicae reared on (A) A. thaliana Col-0 or (B) bak1-5 mutants plants stably expressing double-stranded (ds) GFP (dsGFP), Rack1 (dsRack1), MpOS-D1 (dsMpOS-D1) or Mp10 (dsMp10). Gene expression levels were measured by quantitative reverse transcriptase PCR (qRT-PCR) using specific primers for each aphid gene. Bars show the means ± SE of normalized gene expression levels relative to the dsGFP control, which was set at 1, of three independent biological replicates (n=5 aphids per repeat). Asterisks indicate significant downregulation of gene expression compared to the aphids on dsGFP plants (Student’s t-probabilities calculated within GLM at P <0.05). (C) Fecundity assays of aphids reared on dsRNA transgenic A. thaliana wild type and bak1-5 plants as shown in A, B. Bars represent the mean number of nymphs per plant produced ± SE in 4 independent experiments (n=5 plants per experiment). Asterisks indicate significant difference compared to dsGFP control aphids (Student's t-probabilities calculated within GLM at P<0.05).

**Figure 3:**
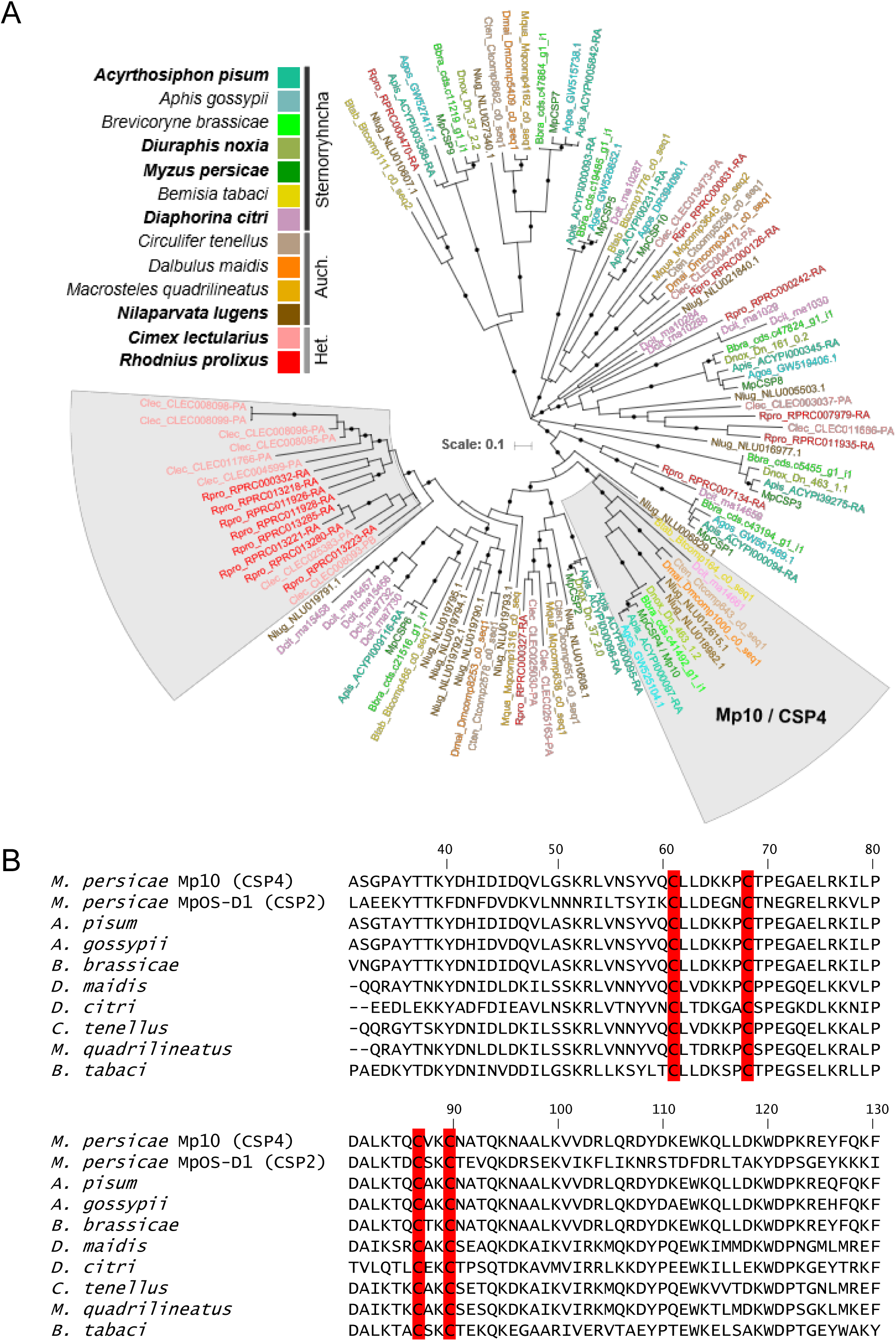
Mp10 homologs are present in diverse plant-feeding insects species of the order Hemiptera and group together as a monophyletic clade. (A) Maximum likelihood phylogenetic tree of CSPs in insect species of the order Hemiptera. Species names at the branches of the tree are colour coded as shown in the upper left legend with species for which whole genome sequence is available shown in bold texts. The tree is arbitrarily rooted at the mid-point, circles on braches indicate SH-like support values greater than 0.8. The Mp10 / CSP4 clade lacking homologs from blood-feeding hemipterans is highlighted with a grey background. The clade with CSPs shared among blood-feeding hemipterans only is also highlighted with a grey background. (B) Alignment of the Mp10/CSP4 amino acid sequences of diverse plant-feeding hemipterans. The four cysteines characteristic of CSP proteins are also conserved (highlighted in red). Alignment was created using Clustal Omega (58).

### Mp10 homologs with ROS burst suppression activity are conserved across sap-feeding hemipteran insects

CSPs are conserved across arthropods, raising the possibility that other plant-feeding hemipterans utilise immune-suppressive CSPs like Mp10. To determine if other hemipteran insects also have Mp10 orthologues we conducted phylogenetic analysis of CSPs among insect species (Supplementary file, Table 4A). CSPs were identified in the publically available genomes of the plant-feeding hemipterans *A. pisum* (pea aphid)*, Diuraphis noxia* (Russian wheat aphid), *Nilaparvata lugens* (rice brown planthopper) and the blood-feeding hemipterans *Cimex lectularius* (the bedbug) and *Rhodnius prolixus* (the kissing bug) (Supplementary file, Table 4B). To increase taxonomic breadth, we also identified CSPs in the transcriptome of *Aphis gossypii* (cotton/melon aphid) and generated de novo transcriptome assemblies from RNA-seq data of the plant-feeding hemipterans *Brevicoryne brassicae* (cabbage aphid), *Macrosteles quadrilineatus* (Aster leafhopper), *Circulifer tenellus* (beet leafhopper), *Dalbulus maidis* (corn leafhopper) and *Bemisia tabaci* (tobacco whitefly) (Supplementary file, Tables 2-4). We identified between 4 and 16 (mean = 8.8) CSPs per species. Phylogenetic analysis of all identified CSP sequences showed strong support for a plant-feeder specific ‘Mp10/CSP4 clade’, with putative Mp10 orthologue proteins present in all analyzed plant-feeding species, but not in blood-feeding species (Fig. 3, Supplementary file, Fig. 4 and 5A However, the blood-feeding hemipteran insects show a CSP clade expansion that was not seen in the plant-feeding hemipterans (Fig. 3A).

Assays to test PTI-suppression activities of GFP fusions of the Mp10/CSP4 orthologues found in *A. pisum, A. gossypii, B. tabaci*, *D. maidis* and *C. tenellus* showed that all suppressed the flg22-induced ROS burst in *N. benthamiana* (Fig. 4A; Supplementary file, Fig 6). We also showed that Mp10 and it’s homologs can supress flg22-induced cytoplasmic calcium bursts-another marker of PTI-(Fig. 4B; Supplementary file, Fig 6). These activity assays together with the phylogenetic analyses suggest that Mp10/CSP4 acquired immunosuppressive activity before the divergence of plant-feeding species of the paraphyletic suborders Sternorrhyncha (aphids and whiteflies) and Auchenorrhyncha (leafhoppers).

**Figure 4:**
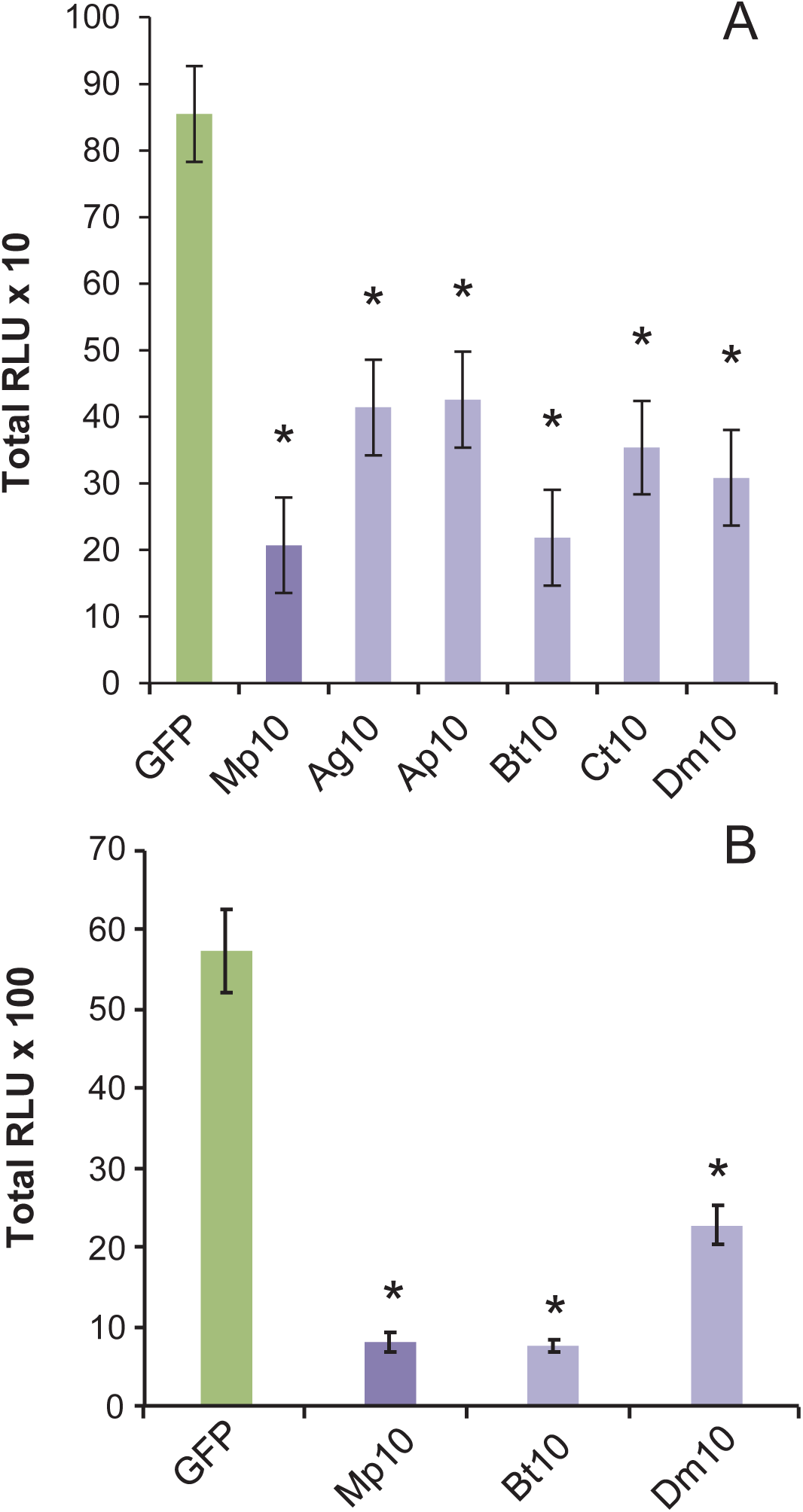
Mp10 homologs from diverse plant-feeding hemipteran species have ROS- and Ca^2+^-burst suppression activities. (A) Total ROS bursts were measured as relative light units (RLU) in luminol-based assays of *N. benthamiana* leaf discs upon elicitation with flg22. (B) Cytosolic calcium bursts measured as RLU using transgenic *N. benthamiana* leaf discs expressing the calcium reporter aequorin. The leaves transiently produced the GFP-tagged homologs of Mp10 from *A. gossypii* (Ag10), *A. pisum* (Ap10), *B. tabaci* (Bt10), *C. tenellus* (Ct10) and *D. maidis* (Dm10) alongside a GFP control. Bars show mean ± SE of total RLUs measured over periods of 60 (A) or 30 (B) minutes after flg22 exposure in three independent experiments (n=8 (A) or n=12 (B) per experiment). Asterisks indicate significant differences to the GFP treatment (Student’s t-probability calculated within GLM at P <0.05).

## Discussion

CSPs are known to regulate behavioural and developmental processes in arthropods. Our findings extend the known functions of CSPs to include the modulation of process in entirely different organisms that are not animals, i.e. in plants. We have demonstrated that proteins belonging to the Mp10 clade from diverse plant feeding hemipteran insects have evolved the ability to act as effector proteins that suppress plant immunity through the suppression of ROS bursts. Strikingly, Mp10 is conserved across two paraphyletic suborders of Hemiptera that diverged over 250 million years ago (21), but absent in blood-feeding hemipteran insects, indicating a key role for this CSP in plant feeding. It is therefore likely that an ancestral CSP4 evolved to suppress plant immunity before the divergence of hemipteran herbivores into two paraphyletic suborders

Importantly, we also demonstrate that Mp10/CSP4 implements its immunosuppressive activity in plants, during aphid feeding, in the plant BAK1 pathway. BAK1 regulates the first layer of the plant defence response and is required for induction of ROS bursts to many pathogens and pests, including aphids (24, 25, 35, 36). We previously detected Mp10 in the cytoplasm and chloroplasts of mesophyll cells located adjacently to aphid piercing-sucking mouthparts (stylets) in aphid feeding sites of leaves (20). Mesophyll cells locate directly below the leaf epidermis and cuticle and are probed by the aphid in the early stages of aphid feeding during navigation of its stylets to the vascular bundle phloem sieve cells, where aphid establish long-term feeding sites (37, 38). Each probe by the aphid involves the delivery of saliva (39). Taken together, these data indicate that Mp10 suppresses ROS bursts in the BAK1-mediated plant defence pathway during the early stages of aphid feeding when these insects introduce saliva into mesophyll cells.

We confirmed earlier reports that overproduction of Mp10 in leaves induces chlorosis (18). It was previously found that the Mp10-induced chlorosis response is dependent on SGT1 (18), a ubiquitin-ligase associated protein that is required for effector-triggered immunity (ETI) (40), which involves recognition of pathogen/pest effectors or effector activities by plant cytoplasmic NBS-LRR resistance proteins leading to cell death or other cellular responses that limit pathogen or pest colonization (41). SGT1 is also required for the induction of chlorosis elicited by the jasmonate isoleucine (JA-Ile) analogue coronatine (produced by the plant-pathogenic *Pseudomonas* species) (42) and the regulation of JA precursor accumulation in chloroplasts upon wounding and herbivory (43). It remains to be seen if Mp10 induces ETI and alters JA accumulation, and if these plant processes and the chlorosis induction are biologically relevant in the context of the aphid-plant interaction, because Mp10 is introduced into a few mesophyll cells and does not seem to move away from the aphid feeding site (20).

We did not find CSP4 clade members in the blood-feeding hemipteran insects *R. prolixus* and *C. lectularius*. However, another CSP clade has expanded specifically in these insects (Fig. 6A). As well, mosquito D7 salivary proteins, which are related to OBPs, prevent collagen-mediated platelet activation and blood clotting in animal host (47–49). The D7 salivary family have expanded in all blood-feeding Diptera, including Culicinae (Culex and Aedes families) and Anopheline mosquitoes, sand fly (Psychodidae) and Culicoides (Family Ceratopogonodae), but are nonetheless quite diverse in sequence (48) that is in line with CSPs and OBPs being prone to birth-and-death evolution and purifying selection (13, 49, 50). Given these attributes and observations that effector genes of plant pathogens are usually fast evolving (51–53), it is surprising that the CSP4 clade has remained conserved in herbivorous hemipteran insects that diverged more than 250 million years ago. Hence, it is highly likely that CSP4 plays a fundamental role in mediating insect-plant interactions.

## Materials and Methods

### Bioinformatics and phylogenetic analyses

To identify *M. persicae* CSPs, published pea and cotton/melon aphid CSPs (13, 14) were BLASTP searched against the GPA clone O genome database. Hits with e<10^−5^ were reciprocally BlastP searched against the *A. pisum* genome, and those with an annotated CSP as the top hit were kept for further analysis. CSPs were also identified in the sequenced genomes of *A. pisum* (pea aphid)*, D. noxia* (Russian wheat aphid), *N. lugens* (rice brown planthopper), *C. lectularius* (the bedbug) and *R. prolixus* (the kissing bug). We sequenced the transcriptomes of *B. brassicae* (cabbage aphid), *C. tenellus* (beet leafhopper), *D. maidis* (corn leafhopper), *M. quadrilliniatus* (aster leafhopper) and *B. tabaci* (tobacco whitefly) using RNAseq (NCBI SRA PRJNA318847-PRJNA318851). Coding sequences (CDS) were predicted from the *de novo* assembled transcriptomes and CSPs were identified in all sets of predicted protein sequences based on reciprocal best blast hits to the annotated set of *M. persicae* CSPs. Annotated CSP sequences from the genome and transcriptome data were aligned and phylogenetic analysis was carried out using this alignment. Full details of the bioinformatics and phylogeneic analysis carried out can be found in SI Appendix, Materials and Methods.

### Cloning

To generate constructs that produce double-stranded RNA, entire coding regions of *Mp10* were introduced into the binary silencing plasmid pJawohl8-RNAi by Gateway cloning technology as described previously (30). For production of N-terminal GFP-fusions, coding regions of *Mp10* and other CSPs without sequences corresponding to signal peptides were cloned into the binary plasmid pB7WGF2 using Gateway technology (54). All constructs were introduced into *Agrobacterium tumefaciens* strain GV3101 containing the helper plasmid pMP90RK.

### Transgenic *A. thaliana* lines

*A. thalian*a (Col-0) ds*GFP* and ds*Rack1* lines generated in Pitino et al. 2011 (30) were used. These lines were crossed with *A. thalian*a (Col-0) *bak1-5* lines as described previously (25, 29). The pJawohl8-RNAi constructs for ds*Mp10* were transformed into *A. thaliana* ecotype Col-0 and the *bak1-5* mutant using the floral dip method, homozygous lines were selected as previously described (30) and then screened for their ability to silence the aphid target genes.

### Plant leaf infiltration and ROS assays

Preparation of agrobacterium for agroinfiltration was carried out as previously described (18). Each construct was infiltrated into the youngest fully expanded leaves of *N. benthamiana* at 4-5 weeks of age, or the leaves of 5-week old Arabidopsis. Leaf discs were harvested 2 (*N. benthamiana)* or 3 (*A. thaliana*) days after infiltration and used in ROS burst assays. Measurements of ROS bursts to 100 nM of the peptide flg22 (QRLSTGSRINSAKDDAAGLQIA;(55); Peptron) and *M. persicae*-derived extract was carried out as previously described (18, 25).

### *M. persicae* survival and fecundity assays

Whole plant survival and fecundity assays were carried out as previously described (56). Each experiment included 5 plants per genotype, and the experiment was repeated on different days to generate data from four independent biological replicates. Five adult aphids from each plant genotype were harvested at the end of the experiment for RNA extraction and use in quantitative real-time PCR analysis.

### Quantitative real-time PCR analysis

RNA extraction, cDNA synthesis and qRT-PCR on aphid samples was carried out as previously described (30). Each sample was represented by the gene of interest and three reference genes (L27, β-tubulin and Actin). Primers are listed in Supplementary Table 5. Mean C_t_ values for each sample-primer pair were calculated from 2 or 3 technical replicates, then converted to relative expression values using (efficiency of primer pair)^−δCt^ (57). The geometric mean of the relative expression values of the reference genes was calculated to produce a normalization factor unique to each sample (58). This normalisation factor was then used to calculate the relative expression values for each gene of interest in each sample. For display of data, mean expression values were rescaled such that aphids fed on ds*GFP* plants represented a value of 1.

### Statistical analysis

All statistical analyses were conducted using Genstat 16 statistical package (VSNi Ltd, Hemel Hempstead, UK). Aphid survival and fecundity assays were analysed by classical linear regression analysis using a Poisson distribution within a generalised linear model (GLM). ROS burst assays and qRT-PCR data were also analysed within a GLM using normal distribution. Means were compared by calculating Student’s t-probabilities within the GLM.

## Acknowledgements

This work was supported by the Biotechnology and Biological Sciences Research Council (BBSRC) grant number BB/J004553/1 awarded to JIC and BBSRC studentships to CD, DP and CW. We thank members of the Hogenhout laboratory at the John Innes Centre (JIC) for materials and useful discussions, Ian Bedford, Gavin Hatt, and Anna Jordan for rearing the insects, the JIC Horticultural Services for taking care of plants, Andrew Davis at JIC for providing photographic services and Cyril Zipfel of the Sainsbury Laboratory, Norwich, for providing flg22 peptides and the *bak1-5* Arabidopsis line.

## Supporting Information Captions

**Figure S1:**
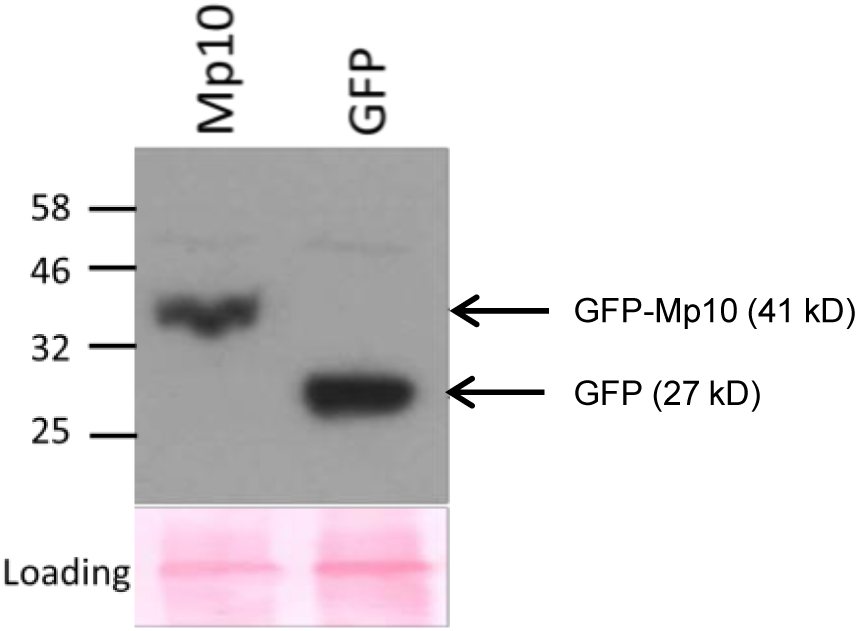
Detection of GFP and GFP-Mp10 proteins in agroinfiltrated *A. thaliana* leaves. The upper panel shows a western blot of protein extracts from four 5 mm diameter leaf discs harvested at 4 days post agroinfiltration from leaves used for ROS assays in Figure 1 A and B. GFP and GFP-tagged Mp10 and MpOS-D1 were detected with antibodies to GFP (arrows at right). The lines at left of the blot indicate the locations of marker proteins in molecular weights (kDa). Protein loading was visualised using Ponceau S solution (loading control in the lower panel).

**Figure S2:**
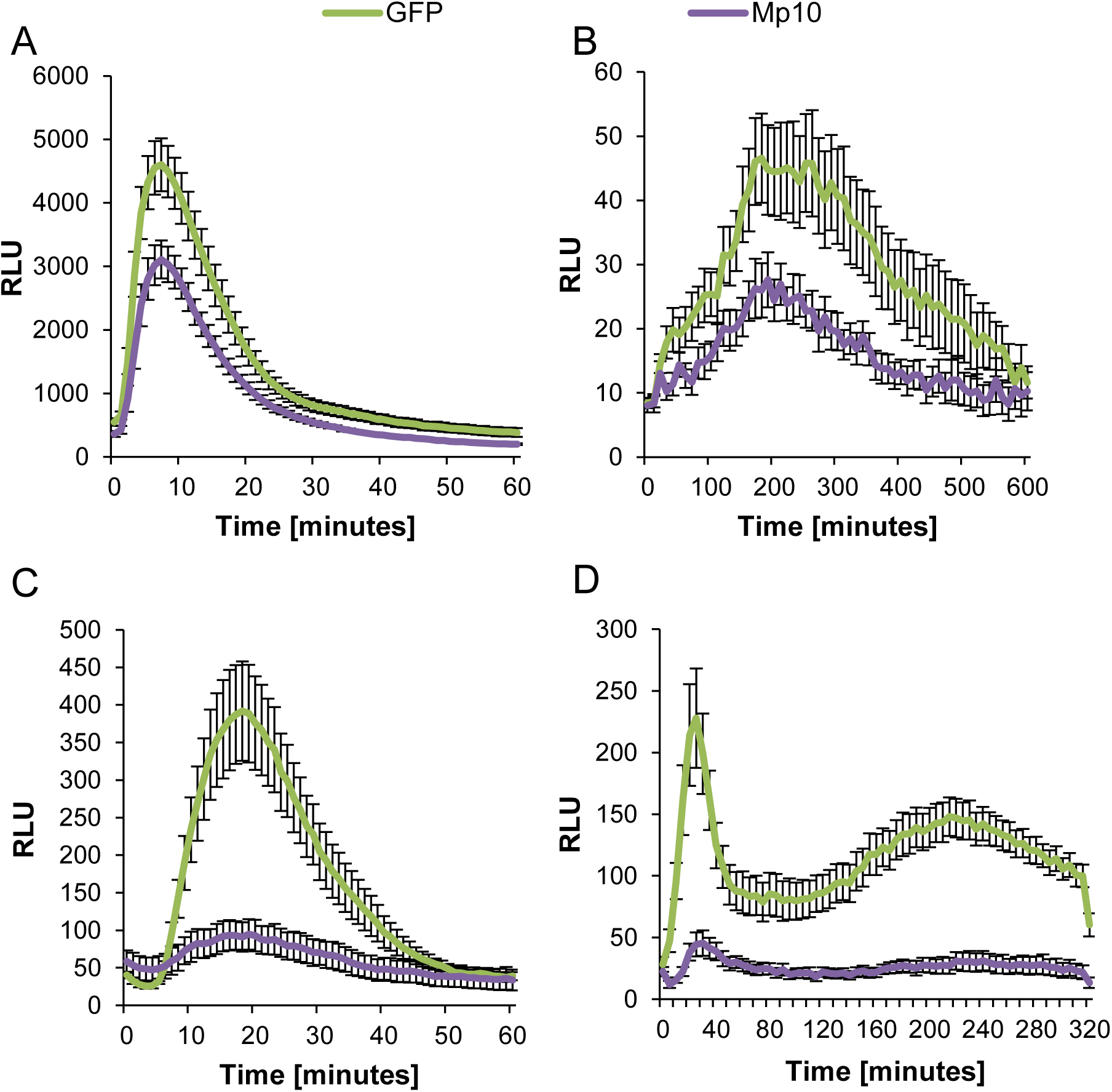
Representative graphs showing Mp10 suppression of elicitor-induced ROS bursts in *A. thaliana* and *N. benthamiana* leaves. Flg22 (A, C) or aphid extract (B, D) were applied to *A. thaliana* (A, B) or *N. benthamiana* (C, D) leaf disks at time point 0. The y-axes show the average ROS bursts in 8 leaf disks measured as relative light units (RLU) in luminol-based assays over 0-60 min (A, C), 0-600 min (B) and 0-320 min (C). The leaves transiently produced GFP-tagged Mp10 (GFP-Mp10) alongside a GFP control.

**Figure S3:**
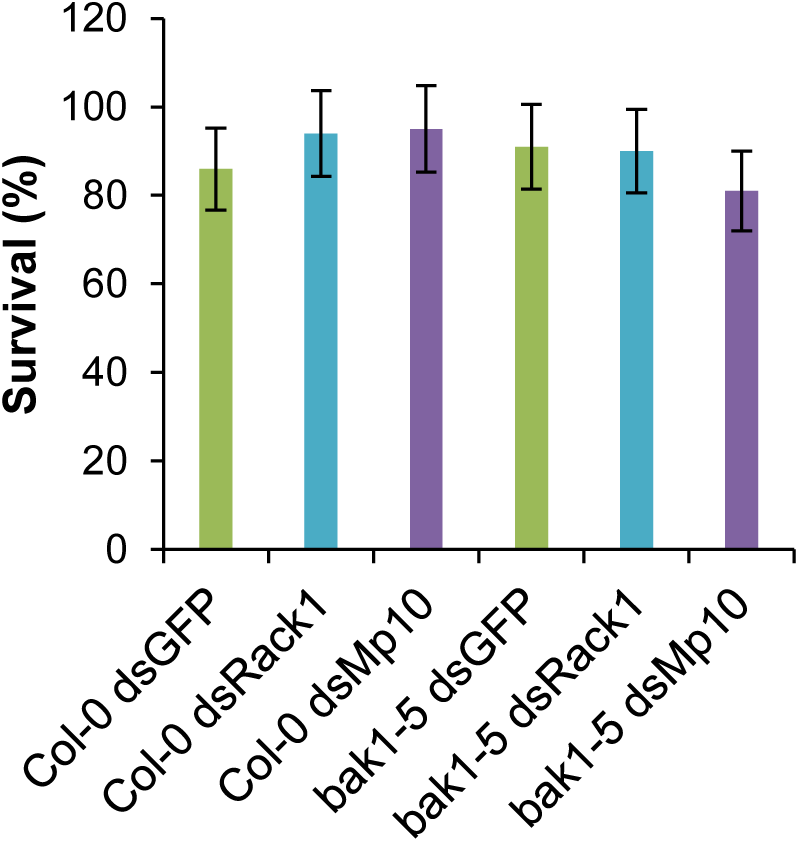
Knock down of *Mp10* expression did not affect aphid survival rates on *A. thaliana* wild type and *bak1-5* mutants. Bars represent the mean number of nymphs alive (out of 5) at the end of the experiment (on day 14, when the final nymph count took place) ± SE in 4 independent experiments (n=5 plants per genotype in each experiment). Asterisks indicate significant difference to *dsGFP* control (Student’s t-probabilities calculated within GLM at P<0.05).

**Figure S4:**
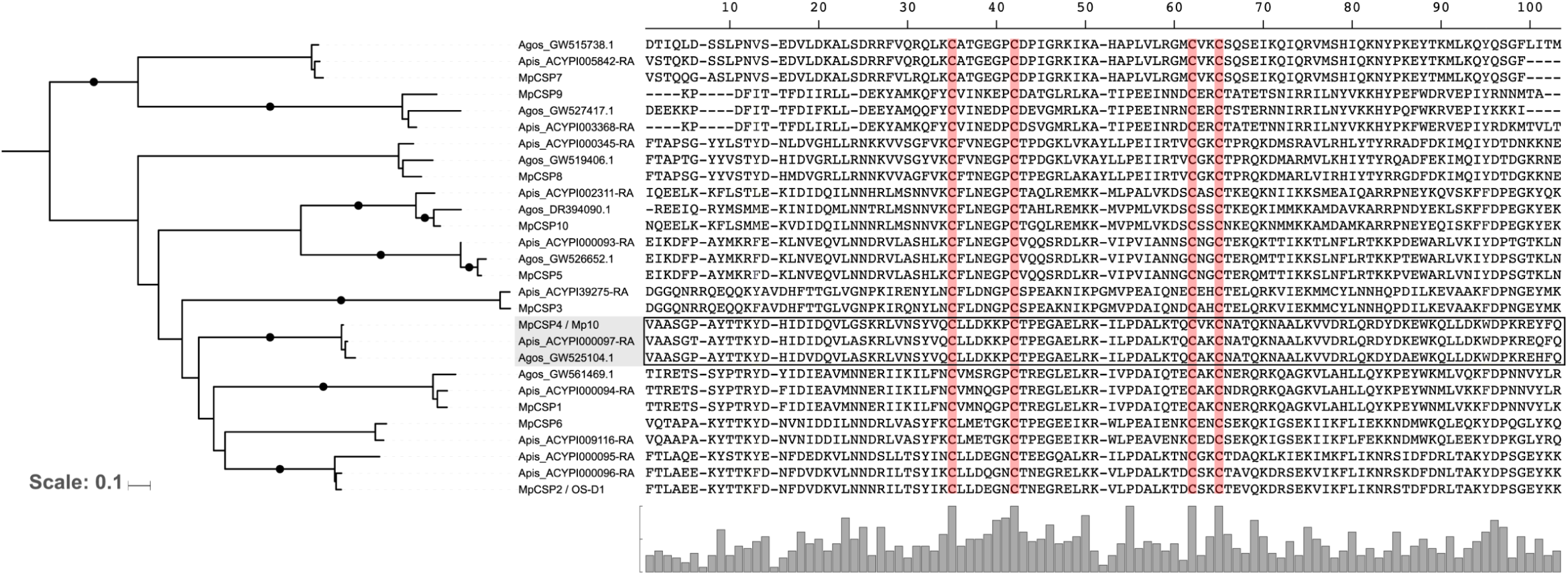
CSPs of the three aphid species *M. persicae*, *A. pisum* and *A. gossypii* cluster into 10 distinct clades including one that contains all Mp10 homologs. Amino acid sequences of CSPs identified in each aphid species were aligned and this alignment was used for generating the phylogenetic tree at left. Conserved cysteines across the CSPs are highlighted in red.

**Figure S5:**
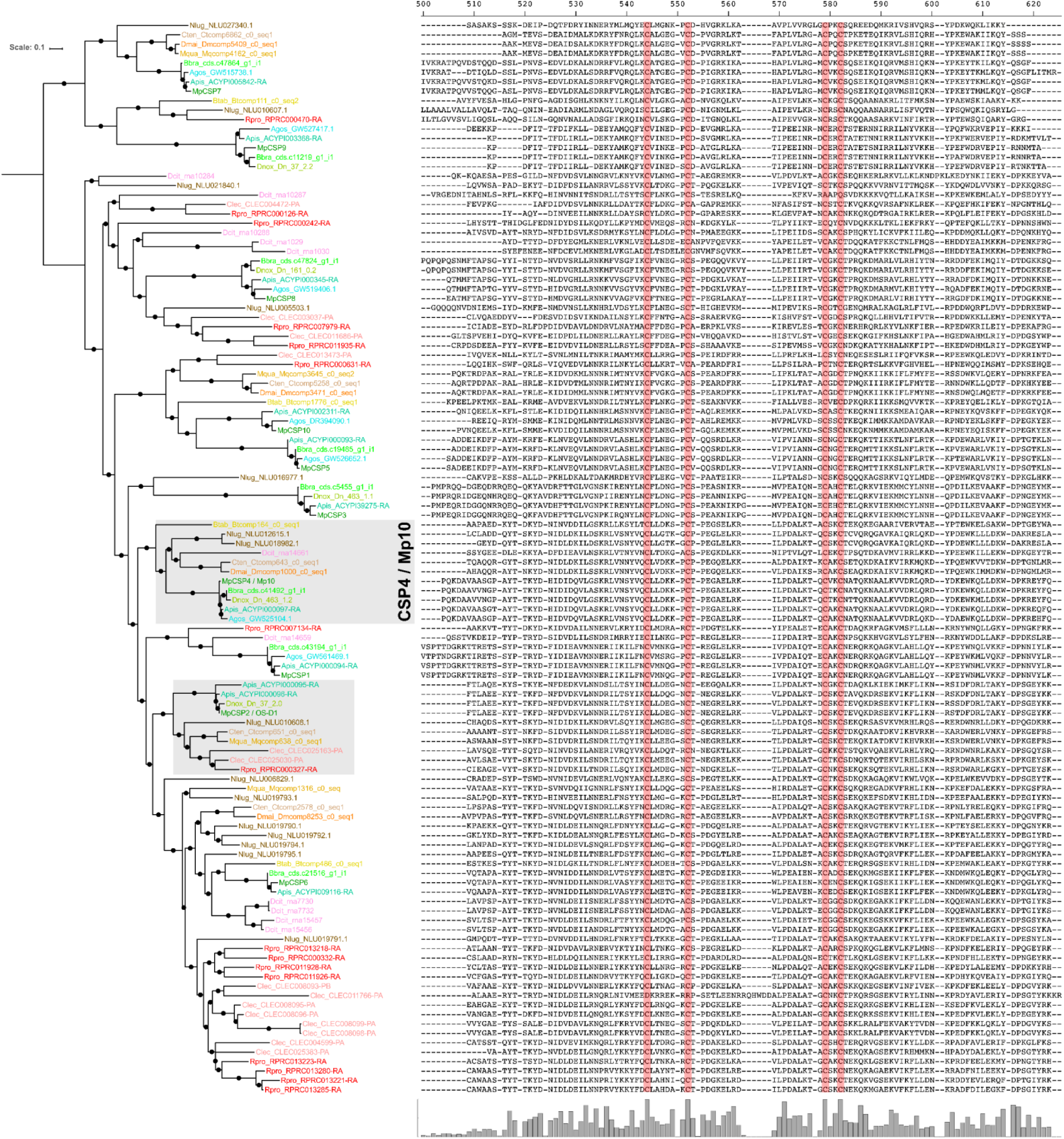
Alignment of CSPs used for building the phylogenetic tree shown in 6A. Amino acid sequences of CSPs identified in available sequence data of insect species in the order Hemiptera were aligned and this alignment was used for generating the phylogenetic trees at left in this figure and for the one shown in Fig. 6A. Conserved cysteines across the CSPs are highlighted in red. The Mp10/CSP4 and clade is highlighted in grey background.

**Figure S6:**
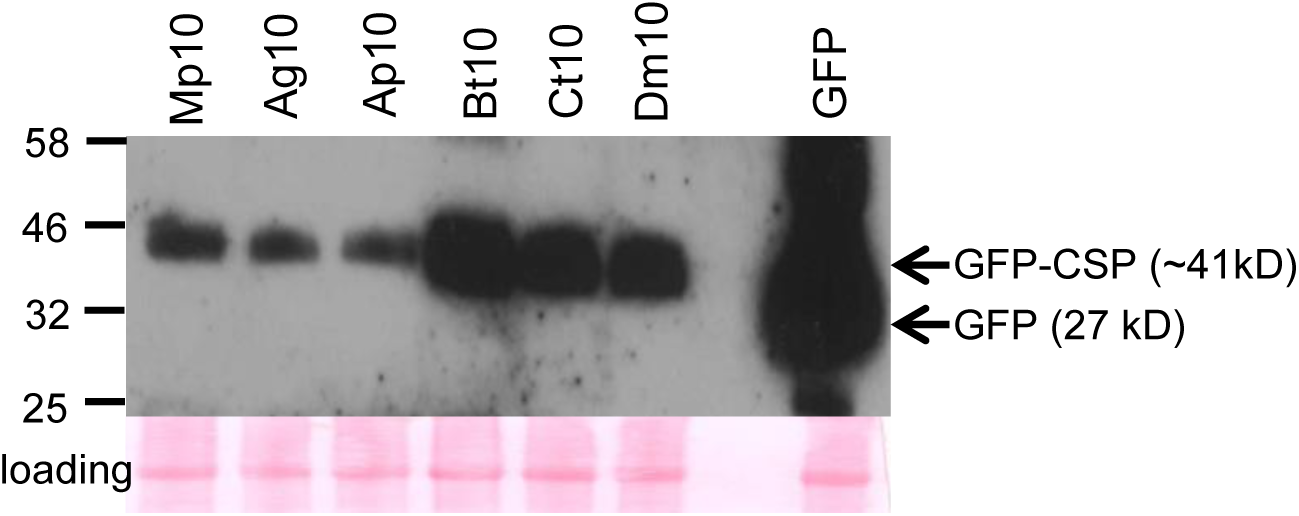
Detection of GFP and GFP fusions of Mp10 homologs from various insect species in agroinfiltrated *N. benthamiana* leaves. The upper panel shows a western blot of protein extracts from two 10 mm diameter leaf discs harvested at 2 days post agroinfiltration from leaves used for ROS assays in Figure 7. The GFP-tagged CSPs were detected with antibodies to GFP (arrows at right). The lines at left of the blot indicate the locations of marker proteins in molecular weights (kDa). Protein loading was visualised using Ponceau S solution (loading control in the lower panel). Abbreviations: Mp, *Myzus persicae*; Ag, *Aphis gossypii*; Ap, A*cyrthosiphon. pisum*; Bt, *Bemisia tabaci*; Ct, *Circulifer tenellus*; Dm, *Dalbulus maidis*.

**Table S1: Blast search statistics for discovery of CSP proteins encoded in the *M. persicae* clone G006 and O genomes.** Query sequence IDs as described in Gu et al (2013(14)).

**Table S2: Sequencing statistics of transcriptomes for five insect species in the order Hemiptera**

**Table S3: Assembly statistics of transcriptome data for five insect species of the order Hemiptera**

**Table S4: CSP annotation.** (A) Summary of hemipteran genomes and transcriptomes searched for CSP sequences. (B) CSP domain annotation for species with sequenced genomes based on hmmer3 searches for PF03392.10. (C) CSPs identified in species with transcriptome data only based on reciprocal best blast hits to *M. persciae* CSPs. (D) summary of filtered CSP sequences retained for phylogenetic analysis.

**Table S5: qRT-PCR primers used in study**

