## Supplemental Table 1 for "Chemosensory proteins in the CSP4 clade evolved as plant immunity suppressors before two suborders of plant-feeding hemipteran insects diverged"

Supplementary Table 1. Blast search statistics for discovery of CSP proteins encoded in the *M. persicae* clone G006 and O genomes. Query sequence IDs as described in Gu et al (2013 (88)).

| Query | Top Hit (G006) | MpCSP ID | Score | E-Value | Similarity | % | Top Hit (O) | Score | E-Value | Similarity | % |
|---|---|---|---|---|---|---|---|---|---|---|---|
| | **Clone G006** | | | | | | **Clone O** | | | | |
| **AGCSP1** | MYZPE13164_G006_v1.0_000014340.1 | MpCSP1 | 273 | 9E-92 | 134/142 | 94 | MYZPE13164_0_v1.0_000025400.1 | 273 | 8E-92 | 134/142 | 94 |
| **AGCSP10** | MYZPE13164_G006_v1.0_000129840.1 | MpCSP10 | 219 | 5E-73 | 122/138 | 88 | MYZPE13164_0_v1.0_000067280.1 | 216 | 5E-72 | 121/137 | 88 |
| **AGCSP2** | MYZPE13164_G006_v1.0_000149850.1 | MpCSP2 / OS-D1 | 234 | 3E-79 | 124/129 | 96 | MYZPE13164_0_v1.0_000099700.1 | 234 | 3E-79 | 124/129 | 96 |
| **AGCSP4** | MYZPE13164_G006_v1.0_000014330.1 | MpCSP4 / Mp10 | 258 | 3E-88 | 139/145 | 96 | MYZPE13164_0_v1.0_000025390.1 | 258 | 3E-88 | 139/145 | 96 |
| **AGCSP5** | MYZPE13164_G006_v1.0_000002320.1 | MpCSP5 | 239 | 3E-81 | 133/139 | 96 | MYZPE13164_0_v1.0_000049890.1 | 241 | 6E-82 | 134/139 | 96 |
| **AGCSP6** | MYZPE13164_G006_v1.0_000014360.1 | MpCSP6 | 243 | 4E-83 | 125/131 | 95 | MYZPE13164_0_v1.0_000025420.1 | 243 | 4E-83 | 125/131 | 95 |
| **AGCSP7** | MYZPE13164_G006_v1.0_000196370.1 | MpCSP7 | 280 | 5E-97 | 142/155 | 92 | MYZPE13164_0_v1.0_000067300.1 | 280 | 5E-97 | 142/155 | 92 |
| **AGCSP8** | MYZPE13164_G006_v1.0_000002330.1 | MpCSP8 | 254 | 2E-86 | 140/162 | 86 | MYZPE13164_0_v1.0_000049900.1 | 254 | 2E-86 | 140/162 | 86 |
| **AGCSP9** | MYZPE13164_G006_v1.0_000149840.1 | MpCSP9 | 213 | 5E-70 | 128/171 | 75 | MYZPE13164_0_v1.0_000099690.1 | 213 | 5E-70 | 128/171 | 75 |
| **APCSP1** | MYZPE13164_G006_v1.0_000014340.1 | MpCSP1 | 385 | 3E-135 | 212/285 | 74 | MYZPE13164_0_v1.0_000025400.1 | 385 | 3E-135 | 212/285 | 74 |
| **APCSP10** | MYZPE13164_G006_v1.0_000129840.1 | MpCSP10 | 205 | 2E-67 | 112/124 | 90 | MYZPE13164_0_v1.0_000067280.1 | 200 | 1E-65 | 111/124 | 90 |
| **APCSP2** | MYZPE13164_G006_v1.0_000149850.1 | MpCSP2 / OS-D1 | 252 | 2E-86 | 128/131 | 98 | MYZPE13164_0_v1.0_000099700.1 | 252 | 2E-86 | 128/131 | 98 |
| **APCSP3** | MYZPE13164_G006_v1.0_000149850.1 | MpCSP3 | 204 | 8E-68 | 112/123 | 91 | MYZPE13164_0_v1.0_000099700.1 | 204 | 8E-68 | 112/123 | 91 |
| **APCSP4** | MYZPE13164_G006_v1.0_000014330.1 | MpCSP4 / Mp10 | 260 | 3E-89 | 141/145 | 97 | MYZPE13164_0_v1.0_000025390.1 | 260 | 3E-89 | 141/145 | 97 |
| **APCSP5** | MYZPE13164_G006_v1.0_000002320.1 | MpCSP5 | 225 | 8E-76 | 130/139 | 94 | MYZPE13164_0_v1.0_000049890.1 | 227 | 1E-76 | 131/139 | 94 |
| **APCSP6** | MYZPE13164_G006_v1.0_000014360.1 | MpCSP6 | 210 | 4E-70 | 125/131 | 95 | MYZPE13164_0_v1.0_000025420.1 | 210 | 3E-70 | 125/131 | 95 |
| **APCSP7** | MYZPE13164_G006_v1.0_000196370.1 | MpCSP7 | 295 | 2E-102 | 148/155 | 95 | MYZPE13164_0_v1.0_000067300.1 | 295 | 2E-102 | 148/155 | 95 |
| **APCSP8** | MYZPE13164_G006_v1.0_000002330.1 | MpCSP8 | 275 | 9E-95 | 145/163 | 89 | MYZPE13164_0_v1.0_000049900.1 | 275 | 8E-95 | 145/163 | 89 |
| **APCSP9** | MYZPE13164_G006_v1.0_000149840.1 | MpCSP9 | 234 | 4E-78 | 144/167 | 86 | MYZPE13164_0_v1.0_000099690.1 | 234 | 4E-78 | 144/167 | 86 |
