## Supplemental Table 2 for "Chemosensory proteins in the CSP4 clade evolved as plant immunity suppressors before two suborders of plant-feeding hemipteran insects diverged"

Supplementary Table 2. Sequencing statistics of transcriptomes for five insect species in the order Hemiptera.

| Sample | Library | Total read pairs | Total read bases |
|---|---|---|---|
| *B. tabaci* pooled adults | Bt15 | 50678690 | 5118547690 |
| *C. tenellus* pooled adults | CtA9 | 45287556 | 4574043156 |
| *C. tenellus* nymphs | CtN3 | 45325224 | 4577847624 |
| *D. maidis* adult males | DmF7 | 45008074 | 4545815474 |
| *D. maidis* adult females | DmM16 | 43120958 | 4355216758 |
| *D. maidis* nymphs | DmN10 | 44939338 | 4538873138 |
| *M. quadrilineatus* adult females | MqF5 | 50874832 | 5138358032 |
| *M. quadrilineatus* adult males | MqM1 | 47512360 | 4798748360 |
| *M. quadrilineatus* nymphs | MqN4 | 42465778 | 4289043578 |
| *B. brassicae* adults | Bb4 | 100776860 | 10178462860 |

Pooled adults = equal numbers of males and females; the nymphs of *C. tenellus*, *D. maidis* and *M. quadrilineatus* were of mixed age of all sub-adult instars.
