## Supplemental Table 3 for "Chemosensory proteins in the CSP4 clade evolved as plant immunity suppressors before two suborders of plant-feeding hemipteran insects diverged"

Supplementary Table 3. Assembly statistics of transcriptome data for five insect species of the order Hemiptera*.

| Species | total length of contigs | total number of contigs | Max Length | Min Length | N90 | N80 | N70 | N60 | N50 |
|---|---|---|---|---|---|---|---|---|---|
| ***B. tabaci*** | 83758742 | 87239 | 21050 | 201 | 326 | 569 | 955 | 1465 | 2048 |
| ***C. tenellus*** | 84379900 | 107826 | 14655 | 201 | 292 | 443 | 687 | 1004 | 1380 |
| ***D. maidis*** | 96317057 | 105906 | 26318 | 201 | 308 | 527 | 895 | 1369 | 1907 |
| ***M. quadrilineatus*** | 89997669 | 141028 | 16096 | 201 | 260 | 347 | 489 | 696 | 974 |
| ***B. brassicae*** | 98445169 | 111056 | 27540 | 201 | 307 | 493 | 837 | 1327 | 1853 |

*transcriptome data shown in Table S2 were assembled; Transcriptomes of pooled adults, males, females and nymphs were combined.
