## Supplemental Table 5 for "Chemosensory proteins in the CSP4 clade evolved as plant immunity suppressors before two suborders of plant-feeding hemipteran insects diverged"

Supplementary Table 5. qRT-PCR primers used in the study.

| Gene name | Sequence (5' -> 3') |
|---|---|
| **Actin** | F GGTGTCTCACACACAGTGCC |
| | R CGGCGGTGGTGGTGAAGCTG |
| **Tubulin** | F CCATCTAGTGTCGCTGACCA |
| | R GTTCTTGGCGTCGAACATTT |
| **L-27** | F CCGAAAAGCTGTCATAATGAAGAC |
| | R GGTGAAACCTTGTCTACTGTTACATCTTG |
| **Mp10** | F GGTCGGAGCGCCGCAAAAAG |
| | R TTGGAACCCAAAACTTGGTCGATGT |
| **MpOS-D1** | F ACCAACGAAGGCCGAGAATTGAGG |
| | R GGCGGTCAAACGATCAAAGTCAGT |
| **MpRack1** | F GGACGTACCACTCGTCGTTT |
| | R CATGATACCCAATCGCTGTG |
