## Supplementary Materials and Methods for "Chemosensory proteins in the CSP4 clade evolved as plant immunity suppressors before two suborders of plant-feeding hemipteran insects diverged"

**SI Materials and Methods**

**Insect Rearing**

*Myzus persicae* (Rothamsted Research lineage genotype O) (28) were reared on Chinese cabbage (*Brassica rapa*, subspecies *chinensis*), *Acyrthosiphon pisum* were reared on broad bean (*Vicia faba*) and *Aphis gossypii* were reared on cotton (*Gossypium hirsutum*) . The insects were maintained in custom-built acrylic cages located in controlled environment conditions with a 14 hour day (18°C) and a 10-h-night (16°C) photoperiod.

*Dalbulus maidis, Circulifer tenellus* and *Bemisia tabaci* colonies were reared on maize (*Zea mays*), sugar beet (*Beta vulgaris*) and Chinese cabbage respectively. The insects were maintained in custom-built acrylic cages in controlled environment conditions at 22^o^C with a 16 hour day and 8 hour night photoperiod.

**Plant Growth Conditions**

*Arabidopsis thaliana* plants used for experiments were germinated and grown in Scotts Levington F2 compost. Arabidopsis seeds were vernalised for 1 week at 4-6^o^C, then grown in a controlled environment room (CER) with a 10 hour day (90 μmol m^–2^ s^–1^) and a 14 hour night photoperiod and at a constant temperature of 22°C. *Nicotiana benthamiana* plants were germinated on Scotts Levington F1 compost and transferred to Scotts Levington F2 after 12 days. Plants were grown in a CER with a 16 hour day (120 μmol m^–2^ s^–1^) and 8 hour night at a constant temperature of 22^o^C.

**Bioinformatics and phylogenetic analysis**

To annotate all the CSPs present in *M. persicae*, protein sequence databases of the annotated whole genome sequence of GPA clone O and available sequences of the pea aphid *Acyrthosiphon pis*um and cotton/melon aphid *Aphis gossypii* were generated. The published pea and cotton/melon aphid CSPs (52, 78) were BLASTP searched against the clone O genome database at cut-off E-values of e^-5^. Identified putative CSPs were reciprocally BLASTP searched against the pea aphid genome at cut-off E-value of e^-5^ to find additional CSPs and to assess if Mp10 and MpOS-D1 were identified.

CSPs were identified in the sequenced genomes of *A. pisum* (pea aphid)*, D. noxia* (Russian wheat aphid), *N. lugens* (rice brown planthopper), *C. lectularius* (the bedbug) and *R. prolixus* (the kissing bug). For each species, the complete set of annotated proteins was reduced to a single longest transcript per gene and searched for sequences containing the CSP PFAM domain (PF03392) using HMMER3 hmmsearch (79). Sequences containing a partial CSP domain (<80% coverage), multiple CSP domains, or that were shorter than 100 amino acids were removed from downstream analyses.

We sequenced the transcriptomes of insectary reared populations of *B. brassicae* (cabbage aphid), *C. tenellus* (beet leafhopper), *D. maidis* (corn leafhopper), *M. quadrilliniatus* (aster leafhopper) and *B. tabaci* (tobacco whitefly) using RNAseq*.* RNA extractions were performed as described (53), separately for males, females and nymphs of *D. maidis* and *M. quadrilliniatus;* for nymphs and adults of *C. tenellus* and adults of *B. tabaci* and *B. brassicae.* RNAseq and assembly was performed by Macrogen (Seoul, Korea), using 100bp PE Illumina reads. Sequence quality and contamination was assessed using FastQC ( v0.10.0), and reads for each species were assembled using Trinity (v r2011-11-26). Sequencing and assembly statistics can be found in Supplementary Tables S2 and S3. Sequence reads are deposited in the NCBI SRA (accession numbers SUB2085065, SUB2085096, SUB2085095, SUB1473271 and SUB2085094 for *B. brassicae, M. quadrilineatus, D. maidis, B. tabaci* (Biotype B) and *C. tenellus* respectively*)*.

We predicted coding sequences (CDS) from the *de novo* assembled transcriptomes using TRANSDECODER (80). The same analysis was also run on a previously published *A. gossypi* EST based transcriptome (78). CSPs were then identified in all sets of predicted protein sequences (translated from TRANSDECODER predicted CDS) based on reciprocal best blast hits to the annotated set of *M. persicae* CSPs. Any Blast hits with e>10^-5^ were rejected.

All annotated CSP sequences from the genome and transcriptome data that contained complete CSP domains and were at least 100 amino acids in length were aligned with MAFFT-EINSI (81) and ML phylogenetic analysis carried using FASTTREE 2.1 (82) under the JTT model of protein evolution with CAT rate variation. Local branch support was tested using the Shimodaira-Hasegawa test. The resulting phylogeny was visualized using ITOL 3 (83). An Mp10 homolog was identified in the de novo transcriptome assembly of the aster leafhopper *Macrosteles quadrilineatus* (Supplementary file, Table S4C) but was omitted from the phylogenetic analysis due to the predicted peptide sequence being shorter than 100 amino acids.
